# Small non-coding RNA expression in developing mouse nephron progenitor cells

**DOI:** 10.1101/289421

**Authors:** Yu Leng Phua, Andrew Clugston, Kevin Hong Chen, Dennis Kostka, Jacqueline Ho

## Abstract

MicroRNAs (miRNAs) are small non-coding RNAs that are essential for the regulation of gene expression and play critical roles in human health and disease. Here we present comprehensive miRNA profiling data for mouse nephron progenitors, which give rise to most of the cell-types of the nephron, the functional units of the kidney. We describe a miRNA expression profile for nephron progenitors, with 162 miRNAs differentially expressed in nephron progenitors when compared to whole kidney. We also annotated 52, and experimentally validated 4 novel miRNAs, which are expressed in developing kidney. Our data is available as a public resource, so that it can be integrated into future studies and analyzed in the context of other functional and epigenomic data in kidney development. Specifically, it will be useful in the effort to shed light on molecular mechanisms underlying processes essential for normal kidney development, such as nephron progenitor specification, self-renewal and differentiation.

## Background & Summary

MicroRNAs (miRNAs) are small, endogenously synthesized non-coding RNAs (~22 nucleotides long) that are critical in a wide variety of biological processes, where they primarily act to fine-tune gene expression at the post-transcriptional and translational level^1^. Biosynthesis of miRNAs begins in the nucleus, where a primary miRNA (pri-miRNA) transcript is first transcribed by RNA polymerase II^1,2^. Subsequently, the endoribonuclease enzyme Drosha-Dgcr8 complex processes pri-miRNAs into individual hairpin-shaped stem loop precursor miRNAs (pre-miRNAs), which are then exported into the cytoplasm via Exportin5^3–5^. In the final steps of miRNA biogenesis, Dicer processes pre-miRNA into mature miRNA, with the miRNA loaded into the RNA-induced silencing complex (RISC)^6–8^. mRNA 3’UTR target recognition by miRNA is primarily dependent on the miRNA seed sequence located within nucleotide position 2-8 of the miRNA, and complementary Watson-Crick base pairing between the miRNA and mRNA allows the RISC complex to catalyse the process of mRNA degradation and/or translational inhibition^9,10^.

Nephron progenitors are multipotent cells that undergo a mesenchymal to epithelial transition to subsequently differentiate into glomerular podocytes, proximal tubules, loops of Henle and distal tubule in the developing kidney^11^. They also self-renew throughout kidney development, and the number of nephron progenitors is one of the primary determinants of nephron endowment at birth^12^. Since no new nephrons are formed postnatally in humans, low nephron number increases an individual’s risk to develop chronic kidney disease in adulthood^13^. While the role of many protein-coding genes in nephron progenitor self-renewal, maintenance and differentiation is well established, the precise role of small non-coding RNAs remains ill-defined. There are several lines of evidence that point to the importance of miRNAs in kidney development and disease. Mutations in *DROSHA* and *DICER1* have been identified and implicated in several human kidney disorders, including paediatric Wilms Tumour and multilocular cystic renal tumours^14^. Conditional ablation of Dicer-dependent miRNAs in mouse nephron progenitor or ureteric bud lineages during renal development resulted in severe renal hypodysplasia (small kidneys) and collecting duct cyst formation, due to aberrant progenitor apoptosis and attenuated cilium length respectively^15,16^. Also, conditional ablation of Dicer or Drosha in glomerular podocytes and renal stroma results in early postnatal death and a wide variety of renal anomalies including podocyte foot process effacement, marked proteinuria, and glomerular aneurysms^17–21^.

However, more precise characterisation of the role of small RNAs in nephron progenitors during kidney development has been hampered by the lack of comprehensive miRNA expression datasets in this context. Given the biomedical relevance of this system, we have performed high throughput small RNA Sequencing (sRNA-Seq) in three biological replicates of embryonic day 15.5 (E15.5) nephron progenitors and whole kidney (Figure 1). Using an adjusted p-value cutoff of 0.05, we identified a total of 162 miRNAs (5p and 3p strand inclusive) out of 792 detectable miRNAs to be differentially expressed in nephron progenitors when compared to whole kidney. Among the top differentially expressed miRNAs are members of the epithelial-specific *miR-200* family, consisting of *miRs-200a, 200b, 200c, 141* and *429*^22^. Levels of *miR-200* family miRNAs^22^ were significantly lower in nephron progenitors, as might be predicted given that nephron progenitors undergo a mesenchymal to epithelial transition upon differentiation. Furthermore, we uncovered 52 novel miRNA species expressed in the developing kidney. Of these miRNAs, 4 were validated via quantitative real-time PCR (qPCR), with 3 via section *in situ* hybridization. In general, the data resource will be useful for researchers studying miRNA related biology in kidney/nephron development.

**Figure 1.**
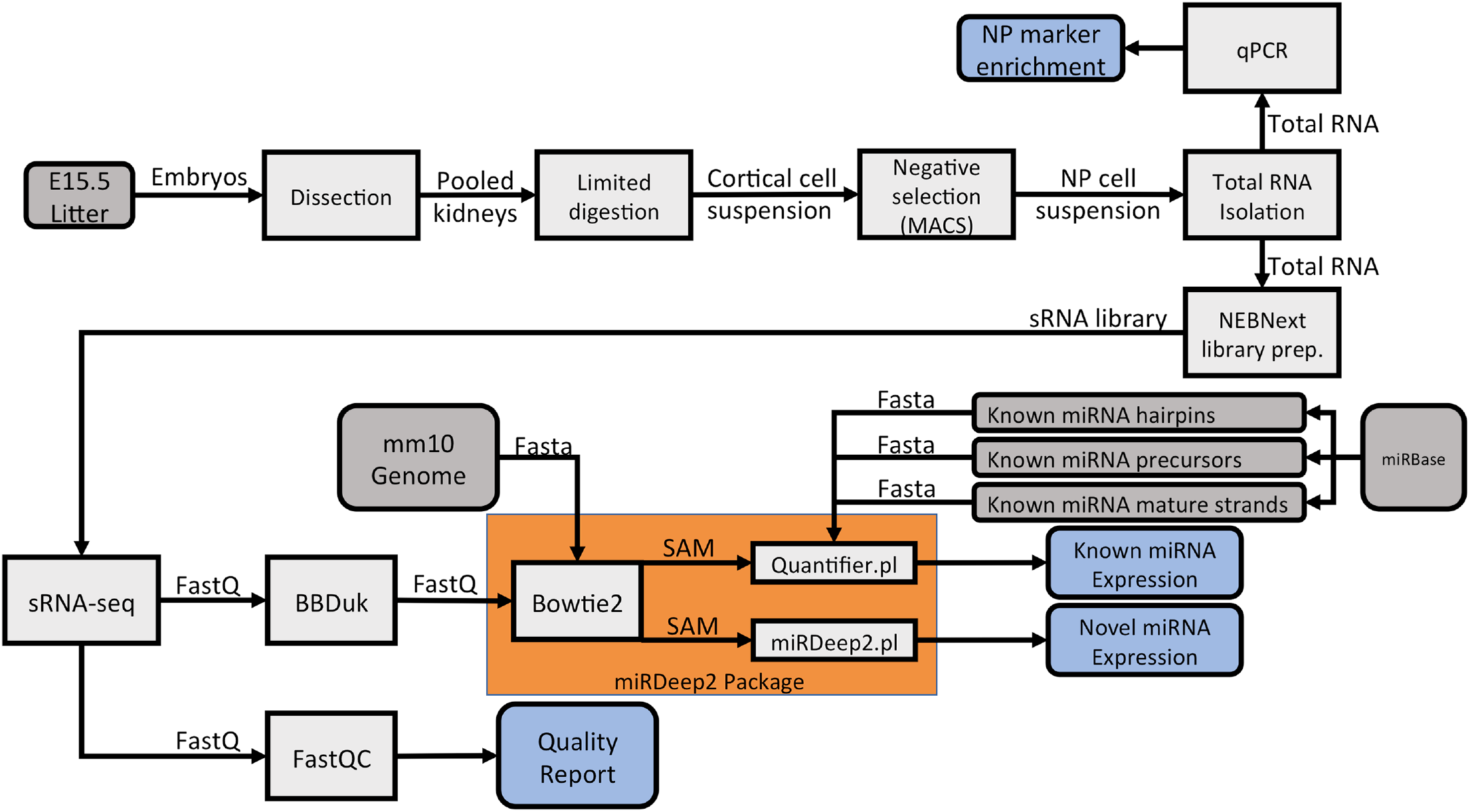
Schematic pipeline illustrating the workflow of nephron progenitor isolation and bioinformatics analysis of the small RNA-Seq dataset. For nephron progenitor isolation, embryonic day 15.5 (E15.5) CD1 mouse kidneys were subjected to limited digestion, followed by negative cell selection through Magnetic Activated Cell Sorting (MACS). Total RNA from the isolate was extracted and subjected to qPCR analysis for enrichment of nephron progenitor markers. Following verification of nephron progenitor enrichment, total RNA was used as an input for the NEBNextMultiplex Small RNA Library Prep to generate libraries for the sRNA-Seq. The sRNA-Seq dataset was analysed in accordance with the pipeline, with the fastq files first analysed by FastQC to determine the quality of the sequencing reads, followed by adaptor removal using the *BBDuk* package, and finally aligned, quantified and annotated to the *mus musculus* mm10 genome using the miRDeep2 package.

## Methods

### Nephron progenitor isolation and total RNA preparation

Nephron progenitors were isolated from 3 litters of E15.5 CD1 mouse embryos (Charles River Laboratories) in accordance to a published protocol using a negative selection approach^23^. Briefly, intact embryonic kidneys were subjected to limited digestion, followed by incubation with a cocktail of monoclonal biotinylated antibodies (eBioscience; CD140a #13-1401-82, CD105 #13-1051-82, Epcam #13-5791-82 and Ter119 #13-5921-82), and magnetic activated cell sorted using Dynabeads MyOne Streptavidin C1 magnetic beads (Thermo; #65001) to deplete unwanted cell types (Figure 1). To minimise undesired gene expression changes, total RNA from nephron progenitors as well as whole kidney samples were immediately processed in QIAzol Lysis Reagent (Qiagen; #79306) and purified using a miRNeasy Micro Kit (Qiagen; #217084). Purified total RNA samples were stored at −80°C until further processing, and freeze thawing of samples was limited to no more than 2 cycles. Quantitative PCR (qPCR) was carried out using standard SYBR Green detection on a BioRad CFX96 Real Time PCR Instrument to determine enrichment of nephron progenitors from the isolation. Primers used include *Six2, Cited1* (nephron progenitors), *Pdgfrβ* (renal stroma), *VE-Cad* (endothelial)*, Calb, Epcam* (epithelial tubules) and *Synpo* (podocytes) (Table 1). Statistical analysis was performed using an unpaired Student’s t-test, and genes with a p-value of <0.05 were considered statistically significant.

**Table 1.**
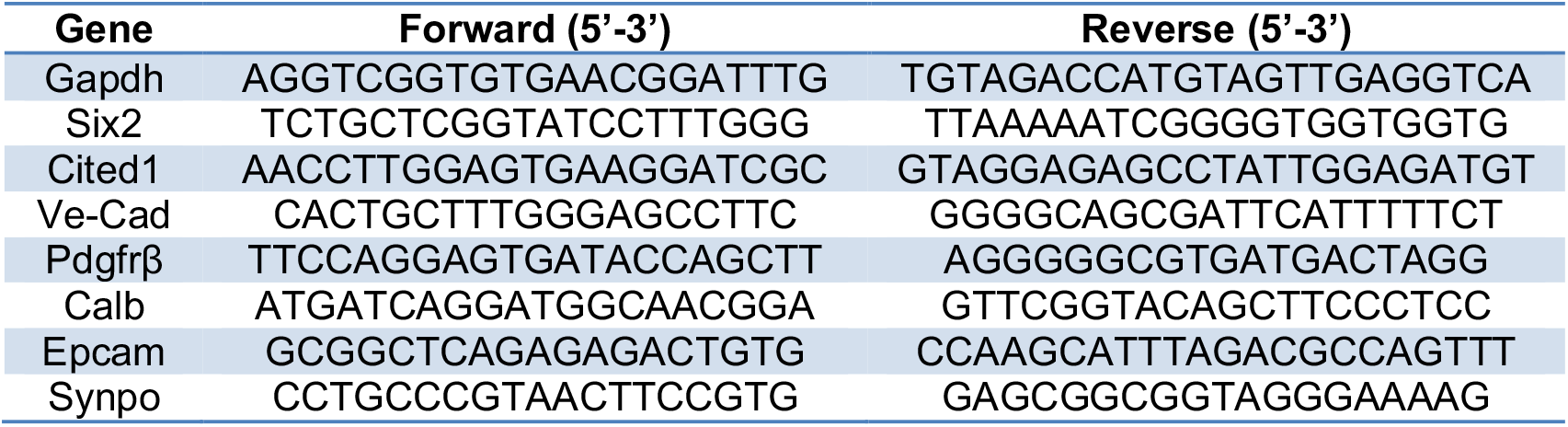
Primer sequences used for quantitative PCR

To account for biological heterogeneity, a total of 3 biological replicates were collected from 3 litters of wildtype CD1 embryos. All experimental procedures were performed in accordance with the University of Pittsburgh Institutional Animal Care and Use Committee guidelines, which adheres to the NIH Guide for the Care and Use of Laboratory Animals. sRNA-Seq experimental workflow and quality control standards were carried out in accordance with ENCODE guidelines (https://www.encodeproject.org/rna-seq/small-rnas/)

### Small RNA library preparation and sequencing

Because miRNAs can be stably bound to mRNAs^24^, total RNA instead of size-selected purified small RNA was used as the initial input for cDNA library synthesis. The NEBNextMultiplex Small RNA Library Prep Set for Illumina (NEB; #E7300S) was used to synthesise barcoded cDNA libraries from 100ng total RNA in accordance with the manufacturer’s protocol and size selection performed using the Agencourt AMPure XP beads (Beckman Coulter; #A63881). The purified cDNA libraries were then pooled, normalised and multiplex sequenced at 1×50bp with the use of a NextSeq 500/550 Mid Output v2 Kit (150 cycles) (Illumina; FC-404-2001) on an Illumina NextSeq 550 System at the Rangos Research Center, yielding approximately 18 million reads per sample.

### Small RNA-Seq data analysis and quality control

Adapter trimming was performed using the BBmap software (https://sourceforge.net/projects/bbmap/) and reads were aligned to the mouse genome (mm10) using Bowtie2^25^. miRNA specific analysis was performed with miRDeep2 software^26^. Expression of known miRNAs (obtained from miRBase v21^27^) was quantified, and miRNAs with more than 10 counts across all sample conditions were subsequently analysed using DESeq2^28^ to identify differential expression between nephron progenitor and whole kidney samples. miRNAs with an Benjamini-Hochberg adjusted p-value of 5% or less are reported. Novel miRNAs were annotated with the standard miRDeep2 parameters, and read alignment data was visualized with the Integrative Genomics Viewer software available from the Broad Institute^29^.

### Data Records

The sRNA-Seq data files are publicly available at NCBI Sequence Read Archive (SRP134975) and Gene Expression Omnibus (GSE111729).

### miRNA Quantitative PCR and Locked Nucleic Acid in situ hybridization validation

cDNA synthesis from total RNA was performed with the use of NCode VILO miRNA cDNA Synthesis Kit (Thermo; #A11193050) in accordance to the manufacturer’s protocol using equivalent amounts of total RNA, and qPCR was performed using standard SYBR Green detection on a BioRad CFX96 Real Time PCR Instrument. The universal reverse primer is supplied with the NCode VILO miRNA cDNA Synthesis Kit, and the forward primer sequence is listed in Table 2. Locked Nucleic Acid miRNA *in situ* hybridisation on E15.5 embryonic kidney cryosections was performed as previously described with the use of custom-designed LNA detection probes^19^ (Table 3) (Exiqon).

**Table 2.**
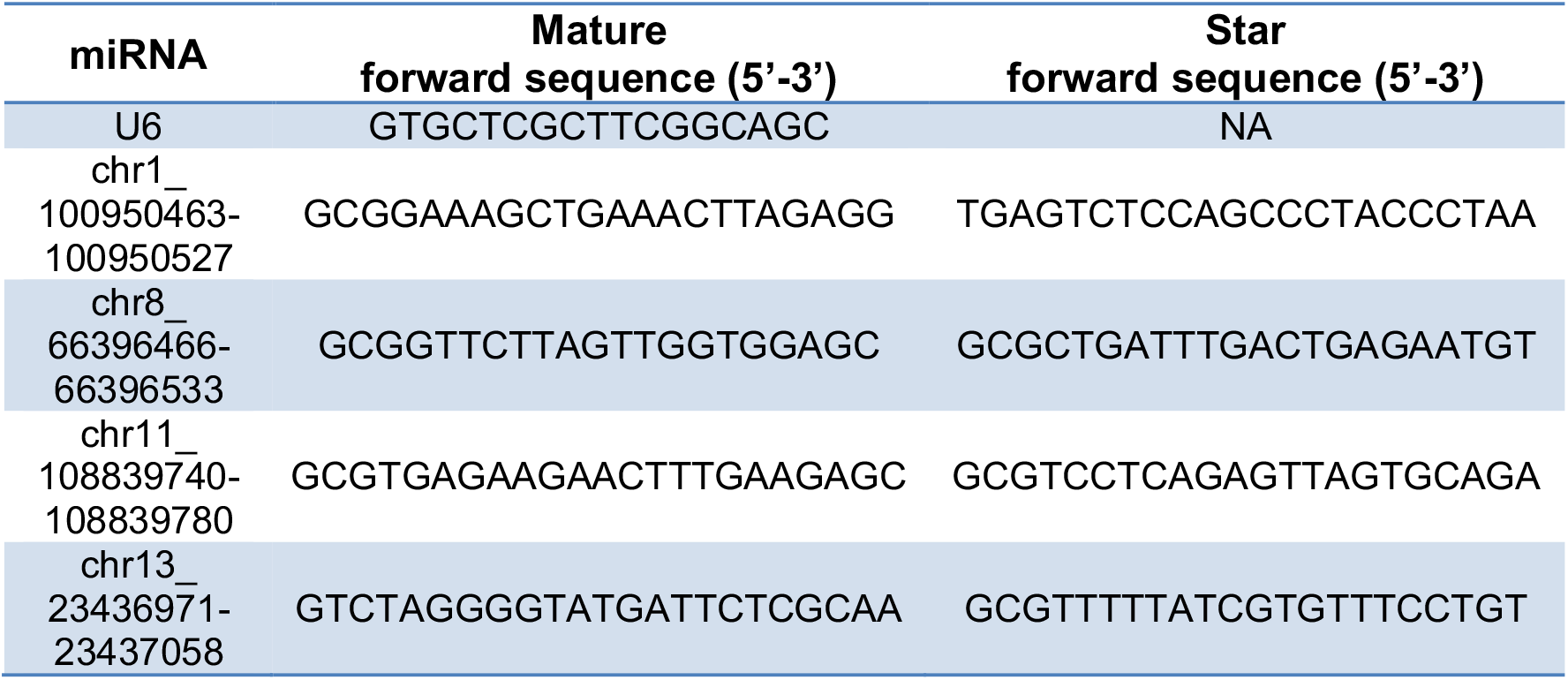
Primer sequences used for novel miRNA quantitative PCR

**Table 3.**
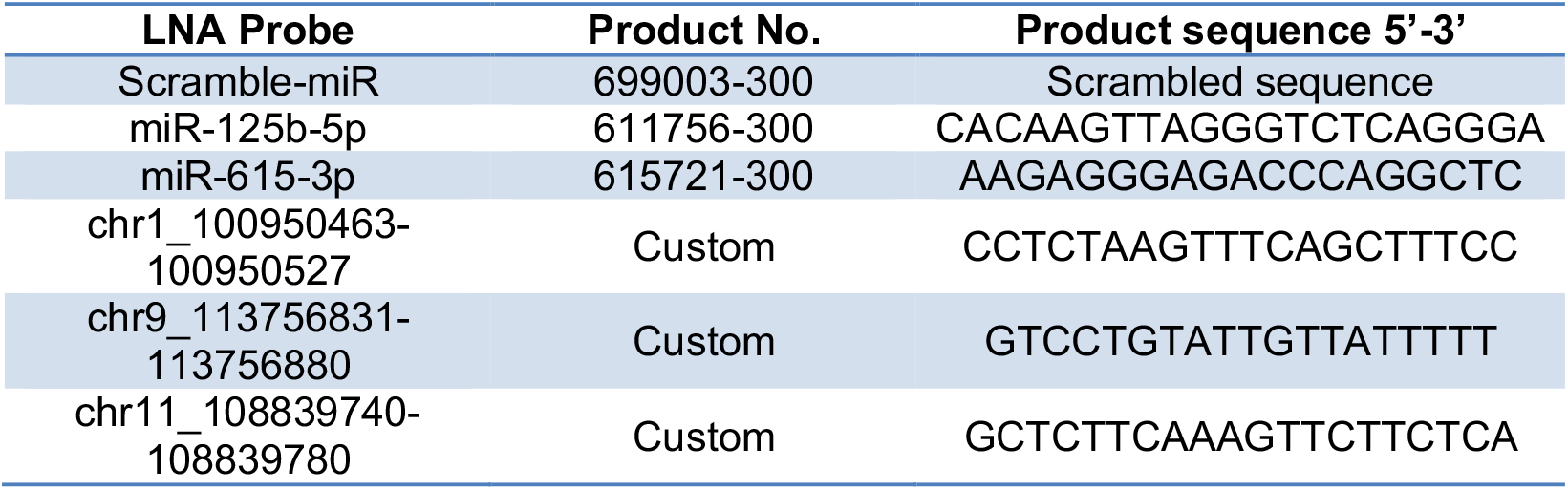
Exiqon miRCURY LNA detection probes used for miRNA section *in situ* hybridisation

## Technical Validation

### Cellular enrichment and total RNA quality control

Our overall goal was to quantify small RNA expression in both mouse nephron progenitors and whole kidney. We used a previously published protocol^23^ to obtain an enriched fraction of nephron progenitor cells from whole embryonic kidney samples. Confirming that this approach was successful, qPCR analysis of the nephron progenitor-enriched isolate showed high enrichment of nephron progenitor transcripts (*Six2*, *Cited1*) in comparison to whole kidney samples, with evidence of minimal cellular contamination from other key renal cell lineages including the endothelium (*VE-Cad*), epithelial tubules (*Calb, Epcam*) and podocytes (*Synpo*) (Figure 2A). An inherent limitation of this protocol^23^ is minor contamination from the renal stroma (*Pdgfrβ*), which was indeed detected in our isolate. Although the Six2-TGC transgenic mouse^33^ provides the opportunity for isolating a more pure fraction of nephron progenitor cells, these mice are known to have renal hypoplasia^34^, suggesting that their nephron progenitors may not be entirely normal. Taken together, our small RNA sequencing dataset represents a highly-enriched fraction of wild-type nephron progenitors, with minor contamination from renal stroma.

**Figure 2.**
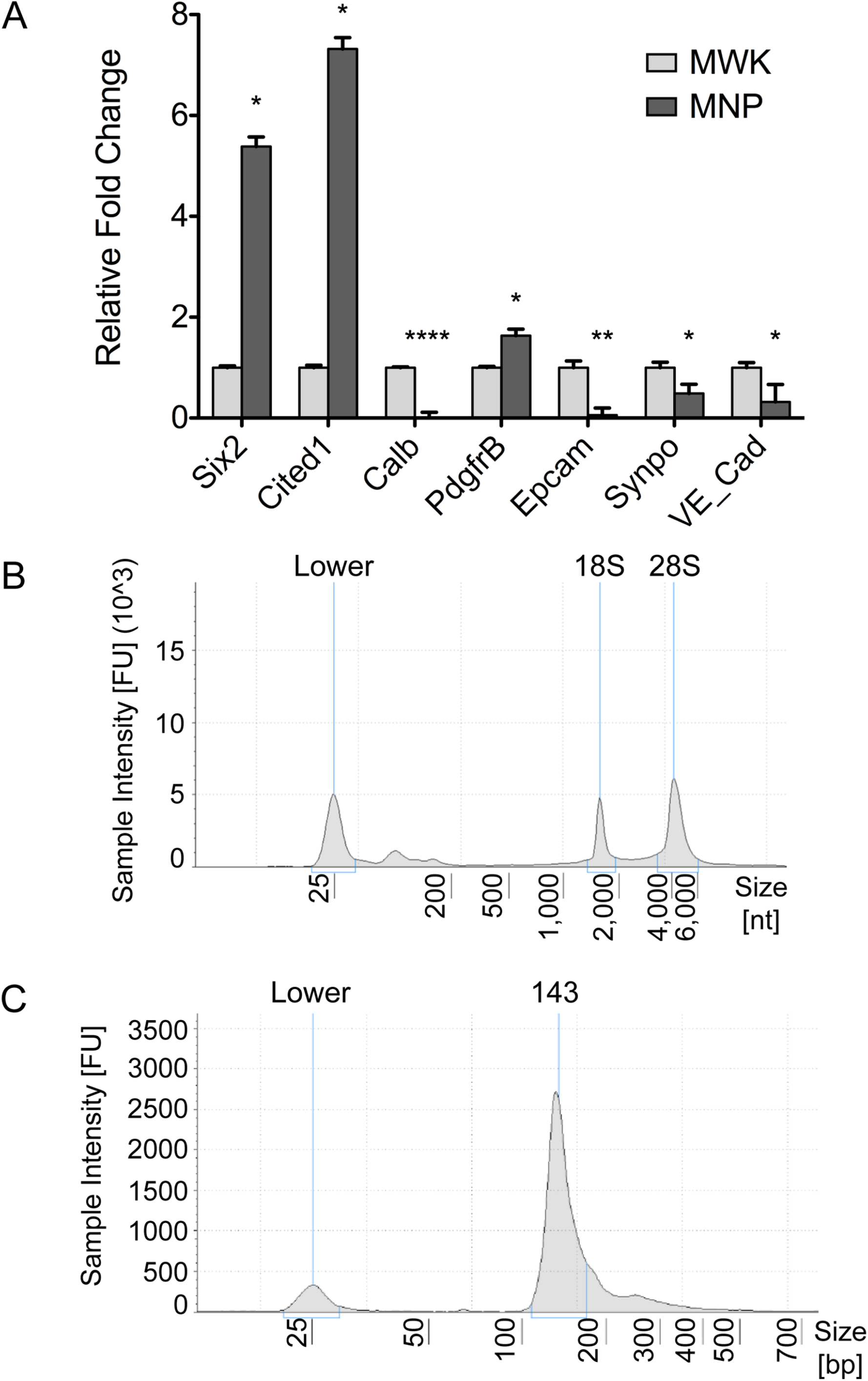
Quality control of the input samples demonstrated an enrichment of nephron progenitors in the cellular isolate with good quality RNA, and successful cDNA library synthesis. (A) qPCR analysis showed that the cells in the isolate are enriched for nephron progenitors (*Six2, Cited1*) with minimal cellular contamination from ureteric bud (*Calb*), podocytes (*Synpo*), endothelial (*VE-Cad*) and epithelial cells (*Epcam*), but minor contamination from the renal stroma lineage (*Pdgfrβ*). (B) TapeStation analysis of the total RNA showed good quality RNA traces with distinct 18S and 28S rRNA peaks with retention of small RNAs as evident by the broad peak to the right of the lower molecular marker. (C) TapeStation analysis showed that the purified cDNA libraries exhibited a distinct peak at ~140-150 nucleotides (adaptor ligated small RNA products), indicating that both the cDNA library construction and cleanup are successful. N=3, * p-value <0.05, ** p-value <0.01, *** p-value <0.001.

Following cellular isolation and total RNA extraction, the quality of the total RNA was assessed on an Agilent 4200 TapeStation System (Agilent; #G2991AA) using a High Sensitivity RNA ScreenTape (Agilent; #5067-5579) to determine the RNA Integrity Number (RIN) score, which is generated based on the electrophoretic profile of 18S and 28S ribosomal RNA (rRNA)^35^ (Figure 2B). All RNA samples used for the sRNA-Seq had a RIN score of above 9 (Table 4) and were therefore considered to be good quality intact RNA with minimal degradation. Retention of small RNA in the total RNA isolation was also evident by a distinct broad peak to the right of the lower molecular marker (Figure 2B). To ensure equivalent amounts of total RNA input for the cDNA library construction, the Qubit RNA BR Assay Kit (Thermo; #Q10210) was used for RNA quantitation.

**Table 4.** Sample IDs and RIN score of RNA samples used for small RNA-Sequencing

| Sample | RIN | Fastq file name |
| --- | --- | --- |
| MNP1 | 10 | MNP1.fastq |
| MNP2 | 10 | MNP2.fastq |
| MNP3 | 10 | MNP3.fastq |
| MWK1 | 9.9 | MWK1.fastq |
| MWK2 | 9.8 | MWK2.fastq |
| MWK3 | 10 | MWK3.fastq |

### Quality control of the cDNA libraries

cDNA libraries were assayed on an Agilent 4200 TapeStation System (Agilent; #G2991AA) using a High Sensitivity D1000 ScreenTape (Agilent; #5067-5584) and were verified to exhibit a distinct peak at ~140-150 nucleotides that corresponds to adapter-ligated constructs derived from small RNAs (Figure 2C). Moreover, TapeStation analysis verified that the final cDNA libraries for sequencing were depleted of large molecular weight products that one would expect be indicative of larger RNA molecules like messenger or ribosomal RNA.

### Quality control of sRNA-Seq data

Purified cDNA libraries for all samples were normalised and multiplex sequenced using a single flow cell on the Illumina NextSeq550 System to minimise technical variability from the sequencing. Following sequencing, reads for each sample was evaluated using the FastQC software (https://www.bioinformatics.babraham.ac.uk/projects/fastqc/) and all sequenced libraries had an average quality score of above 30, indicating high quality base of the sequenced product (Figure 3A). Percentage of aligned reads was consistently above 70%, with 13 million reads mapped per sample on average. For miRDeep2 analysis, aligned reads shorter than 18bp were discarded in keeping with the software’s requirements^26^. Principal component analysis (PCA) revealed a clear distinction between and separation of the nephron progenitor (MNP) and whole kidney (MWK) samples (Figure 3B), as did hierarchical clustering of samples based on their miRNA abundances (Figure 3C). PCA and hierarchical clustering analysis both indicate that samples enriched for nephron progenitors are transcriptionally distinct from whole kidney samples.

**Figure 3.**
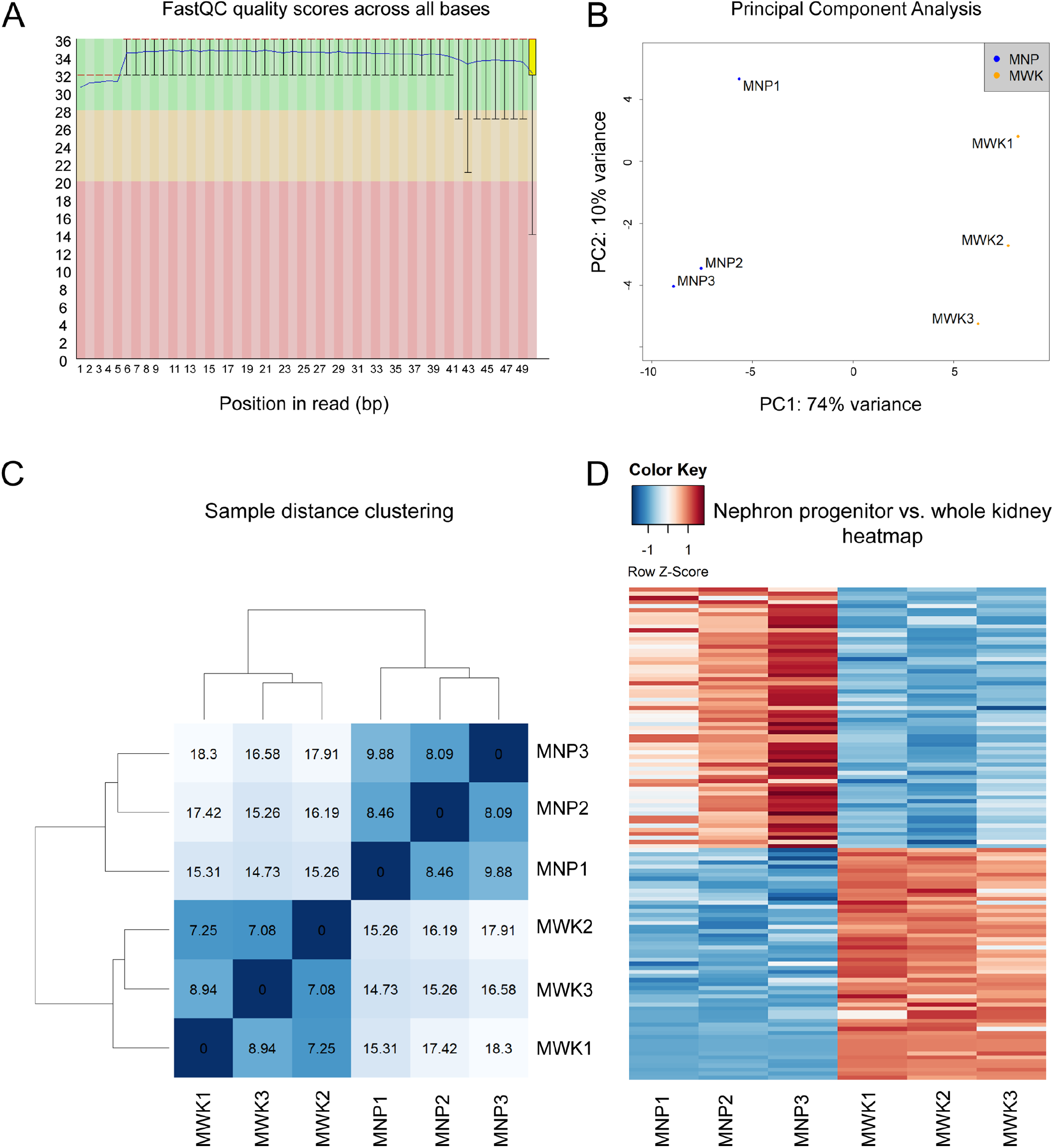
Quality control of the small RNA-Seq dataset showed good sequencing quality reads and congruence of the biological samples. (A) FastQC analysis of a sequenced library (MNP1) showing that each sequenced base had a mean value score of >30, indicating good quality sequencing. (B) Principal Component Analysis of the sRNA-Seq dataset showed clear separation of the nephron progenitor (MNP) and whole kidney (MWK) samples. (C) Hierarchical clustering correctly grouped samples based on their tissue of origin. (D) Heatmap representation of differentially expressed miRNAs in nephron progenitors vs. whole kidney samples with a false discovery rate of 0.05.

We identified a total of 162 miRNAs (5p and 3p strand inclusive) out of 792 detectable miRNAs to be statistically differentially expressed in nephron progenitors relative to whole kidney (Figure 3D, Supplementary Tables 1 and 2). Among the top differentially expressed miRNAs are members of the epithelial-specific *miR-200* family, consisting of *miRs-200a, 200b, 200c, 141* and *429*^22^. Considering that nephron progenitors are mesenchymal in nature and that they undergo a mesenchymal to epithelial transition upon differentiation^33^, the under-expression of the *miR-200* cluster in nephron progenitors not only serves as an excellent internal validation of the phenotypic characteristics of these cells, but also supports the overall integrity of the miRNA profiling dataset.

We explored the use of alternative bioinformatics approaches and obtained comparable results with choices such as *Rsubread* for read mapping to the mm10 genome (parameters: maximum allowed for 3 nucleotides mismatch and 1 insertion-deletion)^36^, *featureCounts* which outputs number of reads assigned to genomic features^37^ and *limma-voom* for differential expression statistical analysis^38^.

### Locked Nucleic Acid section in situ hybridisation and qPCR validation of the small RNA-Seq data

The additional novelty of this dataset stems from its inherent ability to identify novel unannotated miRNA transcripts without *a priori* sequence knowledge. A major challenge in novel miRNA identification is that presumed novel miRNA transcripts are in fact byproducts resulting from degraded mRNA transcripts, and hence not necessarily representative as *bona fide* novel miRNAs. To overcome this, the miRDeep2 package^26^ was used because it includes a series of stringent core analysis modules including the RNAfold tool (http://rna.tbi.univie.ac.at/) and the miRDeep2 core algorithm. The RNAfold tool determines and predicts whether the novel miRNA sequence would exhibit a typical hairpin secondary structure, and the miRDeep2 core algorithm evaluates both the miRNA’s secondary structure and the expected read mapping signature of its hairpin precursor. The core algorithm also checks that the sequencing reads which span the predicted hairpin structure contains specific recognition sites for subsequent processing by Dicer into mature miRNAs. Novel miRNAs were considered high confidence when both mature and star strands were detected in at least 2 independent samples, in conjunction with sequencing reads spanning across the putative chromosome coordinates independently verified on IGV Viewer. In this manner, we identified a total of 52 novel miRNAs, exemplified by the 20 nucleotides long novel miRNA (GGA AAG CTG AAA CTT AGA GG) depicted with a characteristic hairpin precursor secondary structure located within chr1: 100950463-100950527 (Figure 4A and Supplementary Table 3). These novel miRNAs will be submitted for names through miRBase in accordance to their criteria.

**Figure 4.**
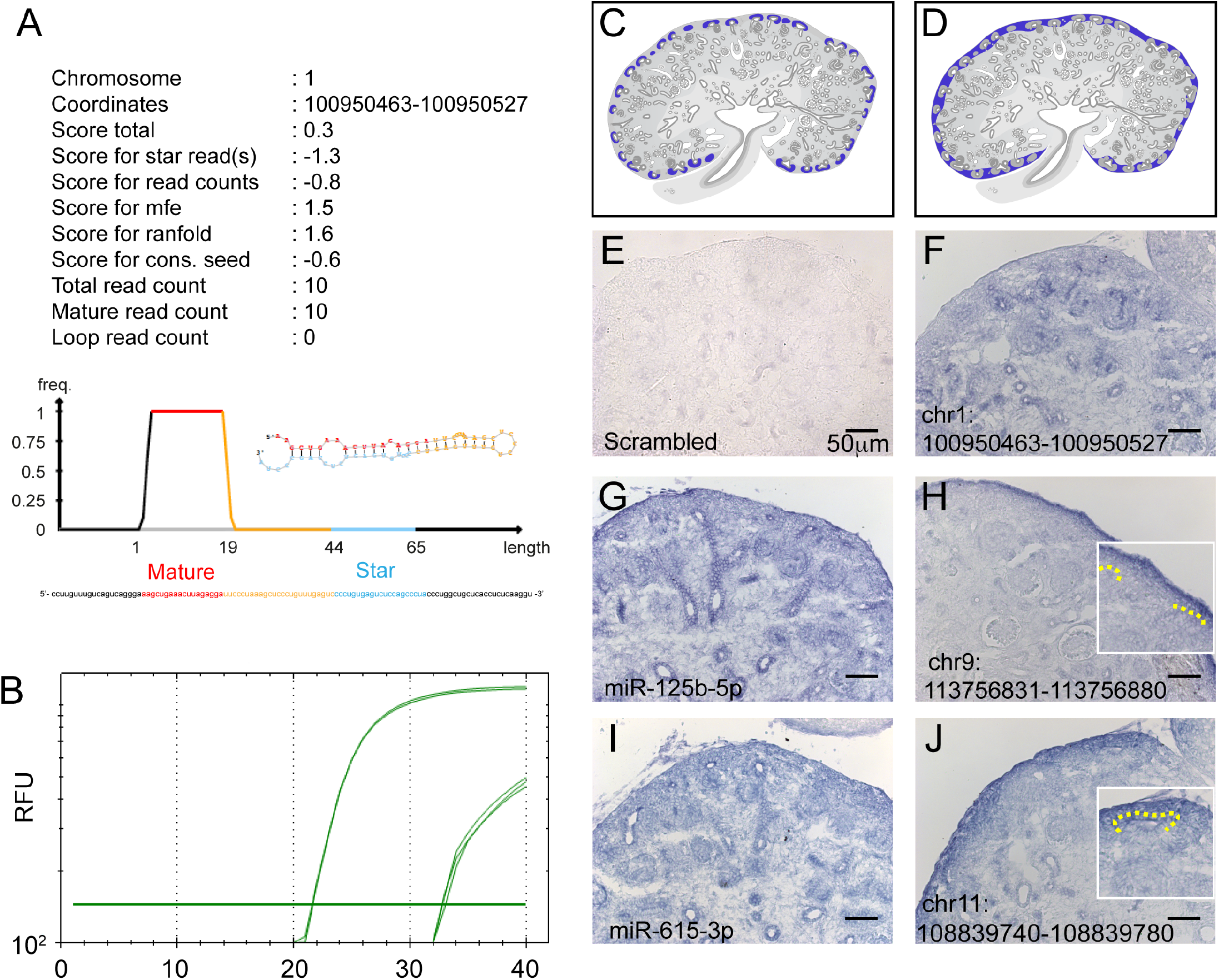
Small RNA-Seq and miRDeep2 analysis revealed novel unannotated miRNAs expressed in the developing kidney. (A) miRDeep2 output for chr1_100950463-10095052, a novel miRNA discovered in our sRNA-Seq data; high frequency (freq) of reads mapped to the mature region of the predicted pre-miRNA structure. (B) qPCR validation of chr1_100950463-100950527 showed a clear distinction between and enrichment of mature (low C_T_) over star strand (High C_T_) transcripts. RFU: Relative Fluorescence Unit. (C and D) For spatial orientation purposes, schematic representative images of nephron progenitors in (C) and renal stroma in (D) are highlighted in blue. (E-J) Locked Nucleic Acid section *in situ* hybridisation was used to validate the spatial expression pattern of miRNAs in the developing kidney. *miRs-125b, 615* and chr1_100950463-100950527 exhibited a ubiquitous expression pattern in the developing kidney. chr9_113756831-113756880 and chr11_108839740-108839780 exhibited a more restricted expression pattern within the nephrogenic zone that comprises both nephron progenitors and renal stroma. Schematics were designed by Kylie Georgas (University of Queensland) and are publicly available for use from the GUDMAP database (http://www.gudmap.org/Schematics).

As a confirmation that the novel miRNA is not an artefact from sRNA sequencing, qPCR was performed on 4 candidate novel miRNAs. The qPCR analysis depicted distinct detection of mature (low C_T_) over star strand (high C_T_) transcripts (Figure 4B and Table 5), thereby validating the overall approach in novel miRNA discovery through sRNA-Seq and miRDeep2 identification. Using miRNA Locked Nucleic Acid *in situ* hybridization, both annotated (*miRs-125b, 615*) and novel miRNAs (chr1_100950463-100950527, chr9_113756831-113756880, chr11_108839740-108839780) were found to be expressed in the developing mouse kidney, with chr9_113756831-113756880 and chr11_108839740-108839780 exhibiting a specific expression pattern within the nephrogenic zone that encompasses both nephron progenitors and renal stroma (Figure 4C). Together, these assays validate both the sRNA-Seq dataset and the approach for novel miRNA discovery in the developing mouse kidney.

**Table 5.** Quantitative PCR profile of novel miRNA assayed

| miRNA | Mature C<sub>T</sub> | Star C<sub>T</sub> |
| --- | --- | --- |
| chr1_<br>100950463-100950527 | 20.25 | 29.01 |
| chr8_<br>66396466-66396533 | 7.04 | 37.65 |
| chr11_<br>108839740-108839780 | 26.59 | 34.48 |
| chr13_<br>23436971-23437058 | 20.60 | 30.65 |

## Usage Notes

The sRNA-Seq data presented here represents a comprehensive resource for miRNA expression in developing nephron progenitors and whole kidney. This resource can be utilised to predict regulatory networks downstream of these expressed miRNAs. For example, this dataset can be integrated with other published RNA-Seq data containing transcriptomic information (mRNA and long non-coding RNA) from nephron progenitors and whole kidney for the analysis of miRNA-mRNA interactions using algorithms such as TargetScan^39^ or DIANA-Tools^40^. Such analysis would allow one to uncover potential candidate miRNAs and/or downstream target mRNAs for future studies. For example, our sRNA-Seq dataset supports the expression of the highly conserved *let-7* family members (*let-7a, 7b, 7c, 7d, 7e, 7f, 7g, 7h, 7i, 7j, 7k* and *miR-98*) in the developing kidney with *let-7c, 7d, 7e* being differentially enriched in nephron progenitors compared to whole kidney during development. The pathogenesis of Wilms tumour has been linked to overexpression of Lin28, a negative regulator of *let-7* microprocessing^41^. Wilms tumour is the most common pediatric kidney cancer and arises from a failure of embryonic kidney tissues to terminally differentiate. Thus, de-repression of *let-7* downstream targets including *Hmga2*^42,43^*, Ras*^44^ and *Myc*^45^ oncogenes may be implicated in the malignant transformation of nephron progenitors in Wilms Tumour^41^. This integrative analysis approach allows for future hypothesis testing studies to elucidate the biological function of key miRNAs in nephron progenitors.

Finally, the sRNA-Seq dataset can be complemented with published ChIP-Seq datasets to infer potential relationships between expressed miRNAs and *cis*-regulatory regions in nephron progenitors. The transcription factor Six2 is known to be critical in maintaining self-renewal and has been shown to regulate target genes that are indispensable for maintaining multipotency of these cells during nephrogenesis^46^. By analysing available Six2 ChIP-Seq data (SRA Study: SRP064623; GEO: GSE73867)^47^ with our current sRNA-Seq dataset, it is now possible to curate candidate miRNAs that are potentially regulated by Six2. Two examples are Six2 binding within the promoter regions of *miRs-210* and *125b*, miRNAs that are enriched in nephron progenitors based on this sRNA-Seq dataset (Figure 5). This is an example of how to use this resource to identify putative Six2-regulated miRNAs that may play important roles in nephron progenitor maintenance and differentiation during kidney development.

**Figure 5.**
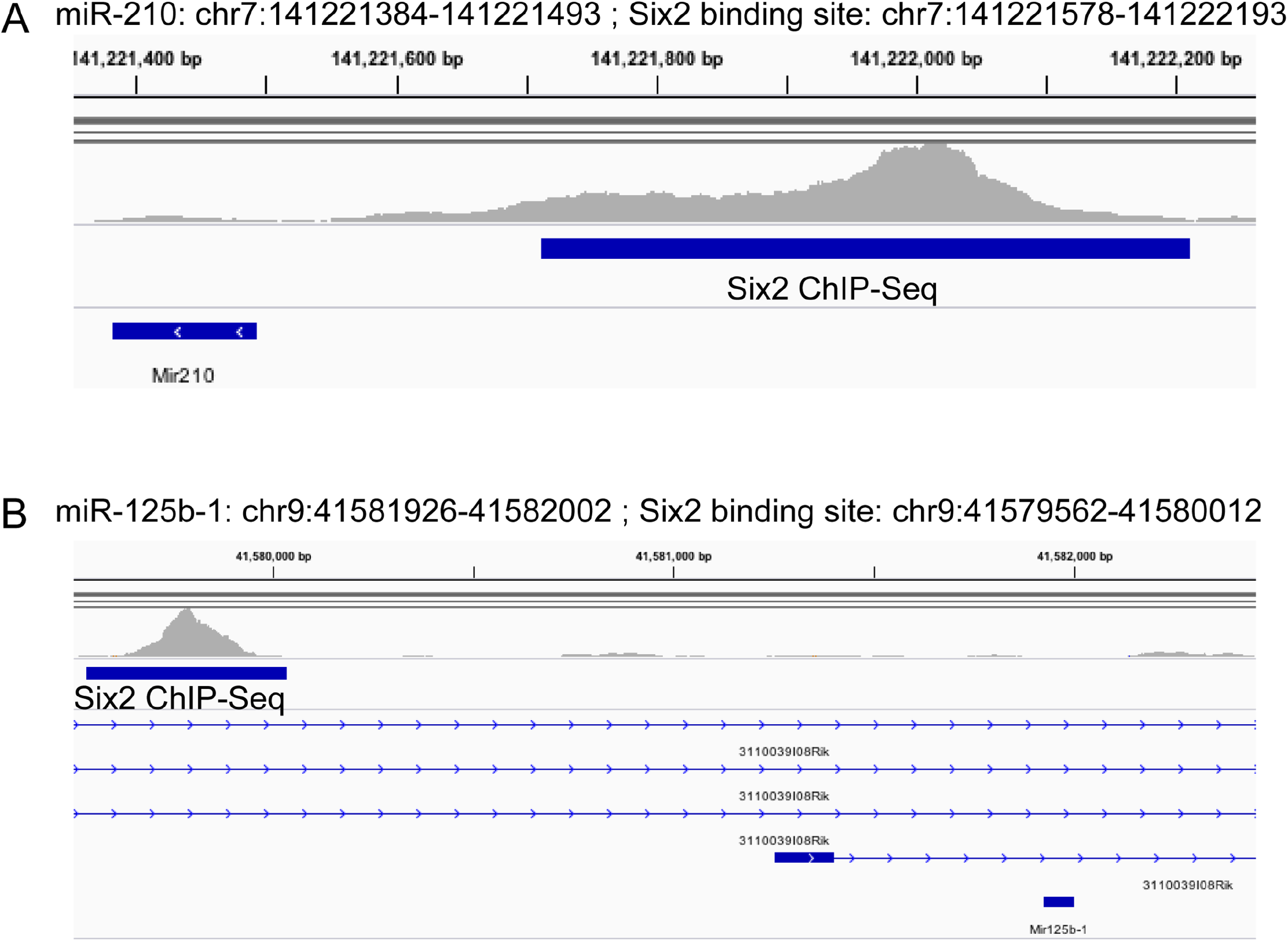
Integrative Genome Viewer visualisation of Six2 binding upstream of expressed miRNAs in nephron progenitors. By combining a publicly available Six2 ChIP-Seq dataset (SRA Study: SRP064623; GEO: GSE73867) with the current sRNA-Seq results, evidence for Six2 binding was observed in the promoter region of *miRs-210* and *125b*, miRNAs that are enriched in nephron progenitors.

## Acknowledgements

Dr. Ho’s laboratory is supported by funding by an NIDDK R00DK087922, NIDDK R01DK103776 and a March of Dimes Basil O’Connor Starter Scholar Award. Dr. Yu Leng Phua was a George B. Rathmann Research Fellow supported by the *American Society of Nephrology* Ben J. Lipps Research Fellowship Program. Kevin Chen was supported by the Research Advisory Committee of Children’s Hospital of Pittsburgh. Dr. Kostka’s laboratory is supported by an NIGMS GM115836. We thank the supercomputing resources provided by the University of Pittsburgh Center for Research Computing.

## Author contributions

YLP and KHC performed the experiments. AC, YLP and KHC analysed the data with supervision from DK and JH. YLP, AC, DK and JH wrote the manuscript, with the final version approved by all authors.

